# Evaluation of the molecular diversity of Brazilian strains of the B.1.1 variant of SARS-CoV-2 used in vaccines

**DOI:** 10.1101/2022.12.23.521847

**Authors:** Iasmin Auta do Nascimento, Lidiane Santos da Silva, Ana Clara da Silva Santos, Pierre Teodósio Felix

## Abstract

In this work, 28 sequences with 57,570 sites of the B.1.1 variant of SARS-CoV-2, from Brazilian states, were used. All sequences (publicly available on the National Center for Biotechnology Information platform (NCBI)) were aligned with Mega X software and all gaps, ambiguous sites and lost data were extracted, resulting in a region in a segment with 8,799 polymorphic (15.2% of the total) that were analyzed for their molecular diversity, F_ST_, demographic and spatial expansion. Phylogenetic relationships of ancestry revealed the absence of genetically distinct subgroups, which was corroborated by the low value of F_ST_ found (15.38%). The low degree of polymorphism found among these samples, corroborated by the almost non-existent genetic distance, helped or established the absence of a genetic structuring pattern, demonstrating a satisfactory pattern of response to vaccines, since all the sequences analyzed were part of the Brazilian strains of variant B.1.1 of SARS-CoV-2 used in vaccines.

## 1. Introduction

With rapid dissemination and with a diversified expansion in specific regions of Brazil, The SARS-CoV-2 establishes itself as the pandemic whose and efforts in the use of protocols, were by far the most inefficient since colonization (Felix *et al*, 2020a).

Diverse factors such as: non-compliance with the health safety protocols and the low population’s compliance to quarantine, increased transmission rates, therefore the number of patients and unfortunately the number of deaths (Felix *et al*, 2020b).

In a very short time, despite the use of vaccines, mutations in the virus have given rise to new variants and subvariants that had a great potential to compromise vaccine-induced immunity (Panke *et al*, 2022). The global spread and increase in infection of the SARS-CoV-2, D614G variant, raised or the question of whether this structural change would compromise the effectiveness of antiviral therapies directed to S protein, especially if they are designed to achieve D614 (Yurkovetskiy *et al*, 2020). The initial subvariant of Ômicron showed from the beginning a strong immunological escape from two doses of the mRNA vaccine, which was only effective with the booster dose (Evans *et al.*, 2022; Gruell *et al.*, 2022). Among the subvariants that emerged from this, the subvariant BA.2 showed a reinfection capacity greater than BA.1, in addition to a slight increase in immunological evasion (Stegger *et al.*, 2022), succeeded by the subvariants BA.4 and BA.5 with identical S proteins, exhibiting greater immunological escape (Kimura *et al.*, 2022; Tuekprakhon *et al*., 2022).

Since we now have Brazilian strains of variant B.1.1 of SARS-CoV-2 used in vaccines, we can try to understand how much variation exists among mutants that still circulate in our country and estimate the risk of immune evasion (if any). Us, from the Laboratory of Population Genetics and Computational Evolutionary Biology (LaBECom-UNIVISA) believe that continuing to contribute to the genetic-population studies of SARS-CoV-2 in Brazil, is still one of the s strategy s for the mitigation of infection, more precisely when trying to establish relationships between the molecular diversity of the virus.

## 2. Objective

Evaluated the molecular diversity of Brazilian strains of variant B. 1.1 of SARS-CoV-2 already used in vaccines.

## 3. Methodology

### Database

The 28 haplotypes of variant B.1.1 of SARS-CoV-2 found in Brazil, were made publicly available and were redeemed from the platform of the National Biotechnology Information Center (NCBI) at the address (https://www.ncbi.nlm.nih.gov/labs/virus/vssi/#/virus?SeqType_s=Nucleotide&VirusLineage_ss=SARS-CoV-2,%20taxid:2697049&Country_s=Brazil&SourceDB_s=GenBank&Lineage_s=B.1.1) on 11 of 10October 2022.

### Phylogenetic analyses

Nucleotide sequences previously described were used for phylogenetic analyses. The sequences were aligned using the MEGA X program (TAMURA *et al.*, 2018) and the gaps were extracted for the construction of phylogenetic trees.

### Genetic Structuring Analyses

Paired F_ST_ estimators, Molecular Variance (AMOVA), Genetic Distance, mismatch, demographic and spatial expansion analyses, molecular diversity and evolutionary divergence time were obtained with the Software Arlequin v. 3.5 (EXCOFFIER *et al.*, 2005) using 1000 random permutations (NEI and KUMAR, 2000). The F_ST_ and geographic distance matrices were not compared. All steps of this process are described below.

#### For Genetic Diversity

Among the routines of LaBECom, this test is used to measure the genetic diversity that is equivalent to the heterozygosity expected in the studied groups. We used for this the standard index of genetic diversity H, described by Nei (1987). Which can also be estimated by the method proposed by PONS and PETIT (1995).

#### For Site Frequency Spectrum (SFS)

According to LaBECom protocols, we used this local frequency spectrum analytical test (SFS), from DNA sequence data that allows us to estimate the demographic parameters of the frequency spectrum. Simulations are made using fastsimcoal2 software, available in http://cmpg.unibe.ch/software/fastsimcoal2/.

#### For Molecular Diversity Indices

Molecular diversity indices are obtained by means of the average number of paired differences, as described by Tajima in 1993, in this test we used sequences that do not fit the model of neutral theory that establishes the existence of a balance between mutation and genetic drift.

#### For Calculating Theta Estimators

Theta population parameters are used in our Laboratory when we want to qualify the genetic diversity of the studied populations. These estimates, classified as Theta Hom – which aim to estimate the expected homozygosity in a population in equilibrium between drift and mutation and the estimates Theta (S) (WATTERSON, 1975), Theta (K) (EWENS, 1972) and Theta (π) (TAJIMA, 1983).

#### For The Calculation of The Distribution of Mismatch

In LaBECom, analyses of the mismatch distribution are always performed relating the observed number of differences between haplotype pairs, trying to define or establish a pattern of population demographic behavior, as described already by (ROGERS; HARPENDING, 1992; Hudson, Hudson, HUDSON, SLATKIN, 1991; RAY *et al*., 2003, EXCOFFIER, 2004).

#### For Pure Demographic Expansion

This model is always used when we intend to estimate the probability of differences observed between two haplotypes not recombined and randomly chosen, this methodology in our laboratory is used when we assume that the expansion, in a haploid population, reached a momentary balance even having passed through τ generations, of sizes 0 N to 1 N. In this case, the probability of observing the S differences between two non-recombined and randomly chosen haplotypes is given by the probability of observing two haplotypes with S differences in this population (Watterson, 1975).

#### For Spatial Expansion

The use of this model in LaBECom is usually indicated if the reach of a population is initially restricted to a very small area, and when one notices signs of a growth of the same, in the same space and over a relatively short time. The resulting population generally becomes subdivided in the sense that individuals tend to mate with geographically close individuals rather than random individuals. To follow the dimensions of spatial expansion, we at LaBECom always take into account:

L: Number of loci

Gamma Correction: This fix is always used when mutation rates do not seem uniform for all sites.

na: Number of substitutions observed between two DNA sequences.
ns: Number of transitions observed between two DNA sequences.
nv: Number of transversions observed between two DNA sequences.
ω: G + C ratio, calculated in all DNA sequences of a given sample.

Paired Difference: Shows the number of loci for which two haplotypes are different.

Percentage difference: This difference is responsible for producing the percentage of loci for which two haplotypes are different.

#### For Haplotypic Inferences

We use these inferences for haplotypic or genotypic data with unknown gametic phase. Following our protocol, inferences are estimated by observing the relationship between haplotype i and xi times its number of copies, generating an estimated frequency (^pi). With genotypic data with unknown gametic phase, the frequencies of haplotypes are estimated by the maximum likelihood method, and can also be estimated using the expected Maximization (MS) algorithm.

#### For The Method of Jukes and Singer

This method, when used in LaBECom, allows estimating a corrected percentage of how different two haplotypes are. This correction allows us to assume that there have been several substitutions per site, since the most recent ancestor of the two studied haplotypes. Here, we also assume a correction for identical replacement rates for all four nucleotides A C, G and T.

#### For Kimura Method with Two Parameters

Much like the previous test, this fix allows for multiple site substitutions, but takes into account different replacement rates between transitions and transversions.

#### For Tamura Method

We at LaBECom understand this method as an extension of the 2-parameter Kimura method, which also allows the estimation of frequencies for different haplotypes. However, transition-transversion relationships as well as general nucleotide frequencies are calculated from the original data.

#### For The Tajima and Nei Method

At this stage, we were also able to produce a corrected percentage of nucleotides for which two haplotypes are different, but this correction is an extension of the Jukes and Cantor method, with the difference of being able to do this from the original data.

#### For Tamura and Nei Model

As in kimura’s models 2 parameters a distance of Tajima and Nei, this correction allows, inferring different rates of transversions and transitions, besides being able to distinguish transition rates between purines and pyrimidines.

#### Minimum Spanning Network

To calculate the distance between OTU (operational taxonomic units) from the paired distance matrix of haplotypes, we used a Minimum Spanning Network (MSN) tree, with a slight modification of the algorithm described in Rohlf (1973). We usually use free software written in Pascal called MINSPNET. EXE running in DOS language, previously available at: http://anthropologie.unige.ch/LGB/software/win/min-span-net/.

#### For Genotypic Data with Unknown Gametic Phase

##### EM algorithm

To estimate haplotypic frequencies we used the maximum likelihood model with an algorithm that maximizes the expected values. The use of this algorithm in LaBECom, allows to obtain the maximum likelihood estimates from multilocal data of gamtic phase is unknown (phenotypic data). It is a slightly more complex procedure since it does not allow us to do a simple gene count, since individuals in a population can be heterozygous to more than one locus.

##### ELB algorithm

Very similar to the previous algorithm, ELB attempts to reconstruct the gametic phase (unknown) of multilocal genotypes by adjusting the sizes and locations of neighboring loci to explore some rare recombination.

#### For Neutrality Tests

##### Ewens-Watterson homozygosis test

We use this test in LaBECom for both haploid and diploid data. This test is used only as a way to summarize the distribution of allelic frequency, without taking into account its biological significance. This test is based on the sampling theory of neutral alllinks from Ewens (1972) and tested by Watterson (1978). It is now limited to sample sizes of 2,000 genes or less and 1,000 different alleles (haplotypes) or less. It is still used to test the hypothesis of selective neutrality and population balance against natural selection or the presence of some advantageous alleles.

##### Accurate Ewens-Watterson-Slatkin Test

This test created by Slatikin in 1994 and adapted by himself in 1996. is used in our protocols when we want to compare the probabilities of random samples with those of observed samples.

##### Chakraborty’s test of population amalgamation

This test was proposed by Chakraborty in 1990, serves to calculate the observed probability of a randomly neutral sample with a number of alleles equal to or greater than that observed, it is based on the infinite allele model and sampling theory for neutral Alleles of Ewens (1972).

##### Tajima Selective Neutrality Test

We use this test in our Laboratory when DNA sequences or haplotypes produced by RFLP are short. It is based on the 1989 Tajima test, using the model of infinite sites without recombination. It commutes two estimators using the theta mutation as a parameter.

##### FS FU Test of Selective Neutrality

Also based on the model of infinite sites without recombination, the FU test is suitable for short DNA sequences or haplotypes produced by RFLP. However, in this case, it assesses the observed probability of a randomly neutral sample with a number of alleles equal to or less than the observed value. In this case the estimator used is θ.

#### For Methods That Measure Interpopulation Diversity

##### Genetic structure of the population inferred by analysis molecular variance (AMOVA)

This stage is the most used in the LaBECom protocols because it allows to know the genetic structure of population measuring their variances, this methodology, first defined by Cockerham in 1969 and 1973) and, later adapted by other researchers, is essentially similar to other approaches based on analyses of gene variance, but takes into account the number of mutations haplotypes. When the population group is defined, we can define a particular genetic structure that will be tested, that is, we can create a hierarchical analysis of variance by dividing the total variance into covariance components by being able to measure intra-individual differences, interindividual differences and/or interpopulation allocated differences.

##### Minimum Spanning Network (MSN) among haplotypes

In LaBECom, this tree is generated using the operational taxonomic units (OTU). This tree is calculated from the matrix of paired distances using a modification of the algorithm described in Rohlf (1973).

##### Locus-by-locus AMOVA

We performed this analysis for each locus separately as it is performed at the haplotypic level and the variance components and f statistics are estimated for each locus separately generating in a more global panorama.

##### Paired genetic distances between populations

This is the most present analysis in the work of LaBECom. These generate paired F_ST_ parameters that are always used, extremely reliably, to estimate the short-term genetic distances between the populations studied, in this model a slight algorithmic adaptation is applied to linearize the genetic distance with the time of population divergence (Reynolds *et al*. 1983; Slatkin, 1995).

##### Reynolds Distance (Reynolds *et al*. 1983)

Here we measured how much pairs of fixed N-size haplotypes diverged over t generations, based on F_ST_ indices.

##### Slatkin’s linearized F_ST’s_ (Slatkin 1995)

We used this test in LaBECom when we want to know how much two Haploid populations of N size diverged t generations behind a population of identical size and managed to remain isolated and without migration. This is a demographic model and applies very well to the phylogeography work of our Laboratory.

##### Nei’s average number of differences between populations

In this test we assumed that the relationship between the gross (D) and liquid (AD) number of Nei differences between populations is the increase in genetic distance between populations (Nei and Li, 1979).

##### Relative population sizes: divergence between populations of unequal sizes

We used this method in LaBECom when we want to estimate the time of divergence between populations of equal sizes (Gaggiotti and Excoffier, 2000), assuming that two populations diverged from an ancestral population of N0 size a few t generations in the past, and that they have remained isolated from each other ever since. In this method we assume that even though the sizes of the two child populations are different, the sum of them will always correspond to the size of the ancestral population. The procedure is based on the comparison of intra and inter populational (π’s) divers that have a large variance, which means that for short divergence times, the average diversity found within the population may be higher than that observed among populations. These calculations should therefore be made if the assumptions of a pure fission model are met and if the divergence time is relatively old. The results of this simulation show that this procedure leads to better results than other methods that do not take into account unequal population sizes, especially when the relative sizes of the daughter populations are in fact.

##### Accurate differentiation population tests

We at LaBECom understand that this test is an analog of fisher’s exact test in a 2×2 contingency table extended to a rxk contingency table. It has been described in Raymond and Rousset (1995) and tests the hypothesis of a random distribution of k different haplotypes or genotypes among r populations.

##### Assignment of individual genotypes to populations

Inspired by what had been described in Paetkau *et al* (1995, 1997) and Waser and Strobeck (1998) this method determines the origin of specific individuals, knowing a list of potential source populations and uses the allelic frequencies estimated in each sample from their original constitution.

##### Detection of loci under selection from F-statistics

We use this test when we suspect that natural selection affects genetic diversity among populations. This method was adapted by Cavalli-Sforza in 1996 from a 1973 work by Lewontin and Krakauer.

## 4. Results

### 4.1. General properties of the sequences analyzed

The 28 sequences of the variant B.1.1 of SARS-CoV-2, from several Brazilian states (Paraná, Mato Grosso, Tocantins, Rio de Janeiro, Pernambuco, Alagoas, São Paulo) and some from unidentified Brazilian states, were classified into 7 groups (OP - Alagoas and São Paulo); (OL-Mato Grosso), (OM - Unreported Geolocation); (ON - Rio de Janeiro, Tocantins, São Paulo and Pernambuco); (OL2 - Unreported Geolocation); (MW - Scientific Platform Pasteur-USP (SPPU) São Paulo) and (MZ - Paraná), which corresponded to 7, 2, 10, 6, 1, 1 and 1 haplotypes, respectively. All 57,570 sites, which once aligned all gaps, ambiguous sites and lost data were extracted, resulting in a region with a segment of 8,799 polymorphic (15.2% of the total). Among the sites that varied, only 2% (1321 sites) were revealed to be informative *parsimonium*. This set was used for the construction of a tree of ancestry utilizing the Maximum Likelihood method, based on the information sites of parsimony. Itis possible to understand that the 28 haplotypes comprised distinct subgroups (showing minimal differences between mutants collected between different Brazilian regions) but with phylogenetic relationships that suggest a very low degree of differentiation of the virus for the whole set (Figure 1).

**Figure 1.**
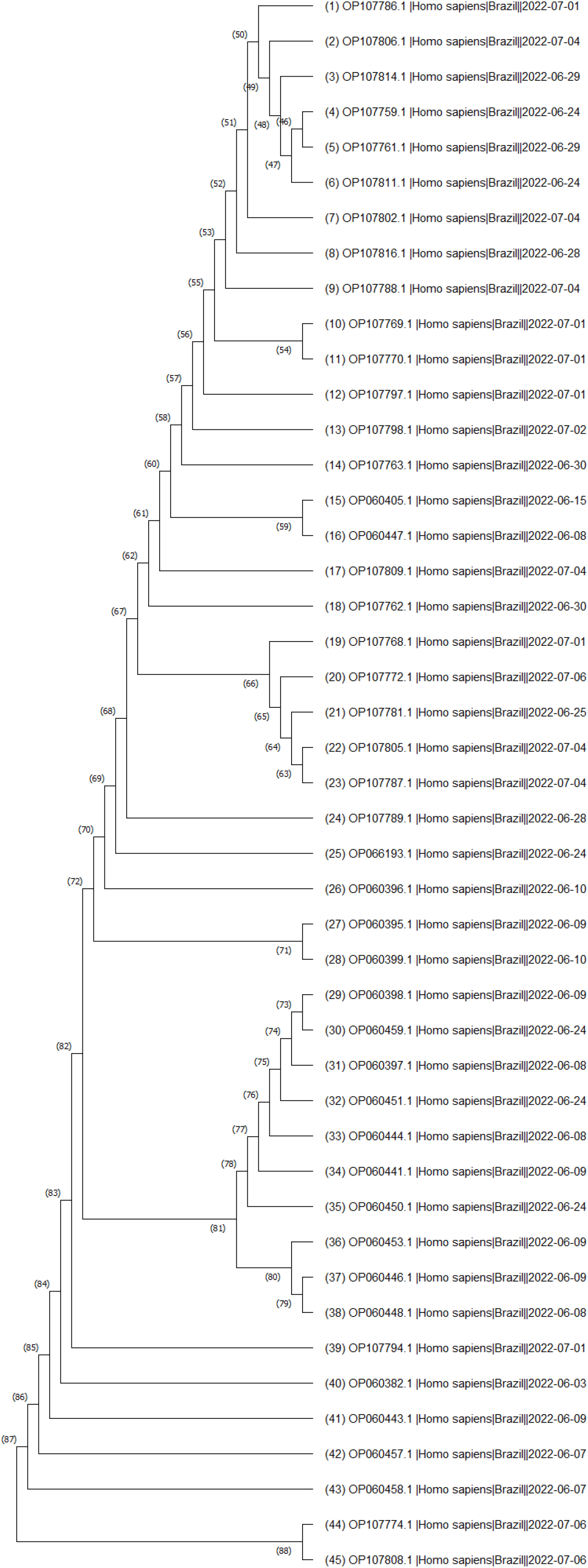
Evolutionary analysis by Maximum Likelihood method. The evolutionary history was inferred by using the Maximum Likelihood method and JTT matrix-based model [1]. The tree with the highest log likelihood (−16188.08) is shown. The percentage of trees in which the associated rate clustered together is shown next to the branches. Initial tree(s) for the heuristic search were obtained automatically by applying Neighbor-Join and BioNJ algorithms to a matrix of pairwise distances estimated using the JTT model, and then selecting the topology with superior likelihood value. A discrete Gamma distribution was used to model evolutionary rate differences between sites (5 categories (+G, parameter = 200.0000)). The rate variation model allowed for some sites to be evolutionarily invariable ([+I], 0.00% sites). This analysis involved 28 sequences. There were a total of 1321 positions in the final dataset. Evolutionary analyses were conducted in MEGA X [2]

1. Jones D.T., Taylor W.R., and Thornton J.M. (1992). The rapid generation of mutation data matrices from protein sequences. Computer Applications in the Biosciences 8: 275-282.

2. Kumar S., Stecher G., Li M., Knyaz C., and Tamura K. (2018). MEGA X: Molecular Evolutionary Genetics Analysis across computing platforms. Molecular Biology and Evolution 35:1547-1549.

### 4.2. Genetic Distance Analysis

Then analyses based on F_ST_ values also confirmed the absence of distinct genetic “entities” with a 5% variation component s within each studied group without significant evolutionary divergences for all samples for a *p-value* of less than 0.05 (Table 1).

**Table 1.**
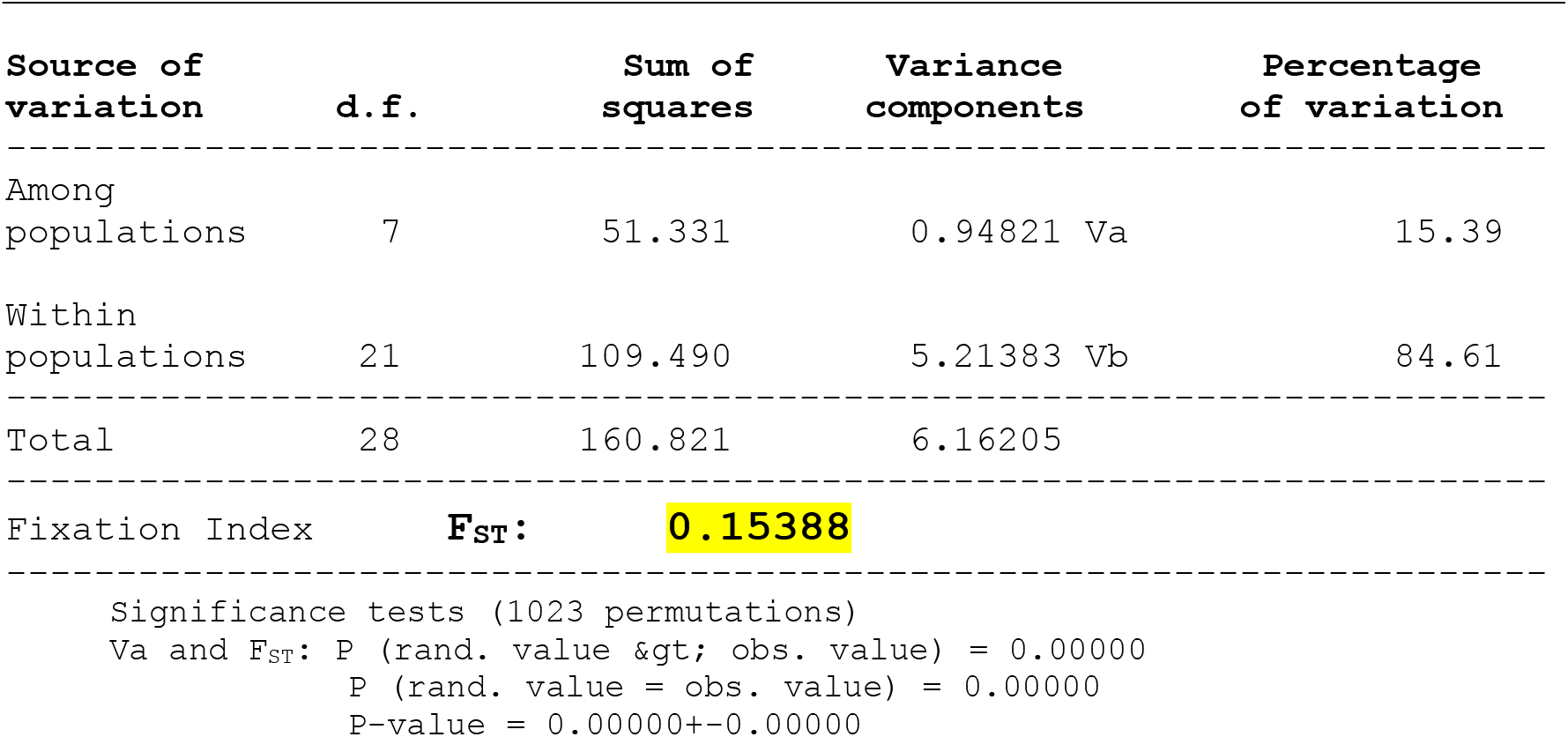
Components of haplotypic variation and paired F_ST_ value for the 28 sequências da variante B.1.1 de SARS-CoV-2, provenientes de Estados brasileiros

Despite the tests for genetic distance do not reveal genetic dissimilarity among all haplotypes, the use of the divergence matrix in the construction of the tree helped in the recognition of minimal similarities between some haplotypes, including for different geographical points. The maximum divergence patterns were also obtained when less robust methods of phylogenetic pairing (e.g., UPGMA) were used, reflecting the non-haplotypic structure in the clades. With the use of a divergence matrix, it was possible to identify geographical variants that had less genetic distances and the probabilities *a posteriori was* able to separate the main clusters into additional small groups, confirming the presence of a minimum probability of kinship between haplotypes. The *Tau* variations (9%), related to the groups, revealing very limited time of divergence, supported by mismatch analysis and demographic and spatial expansion analyses (Figure 2, Figure 3, Table 2).

**Figure 2.**
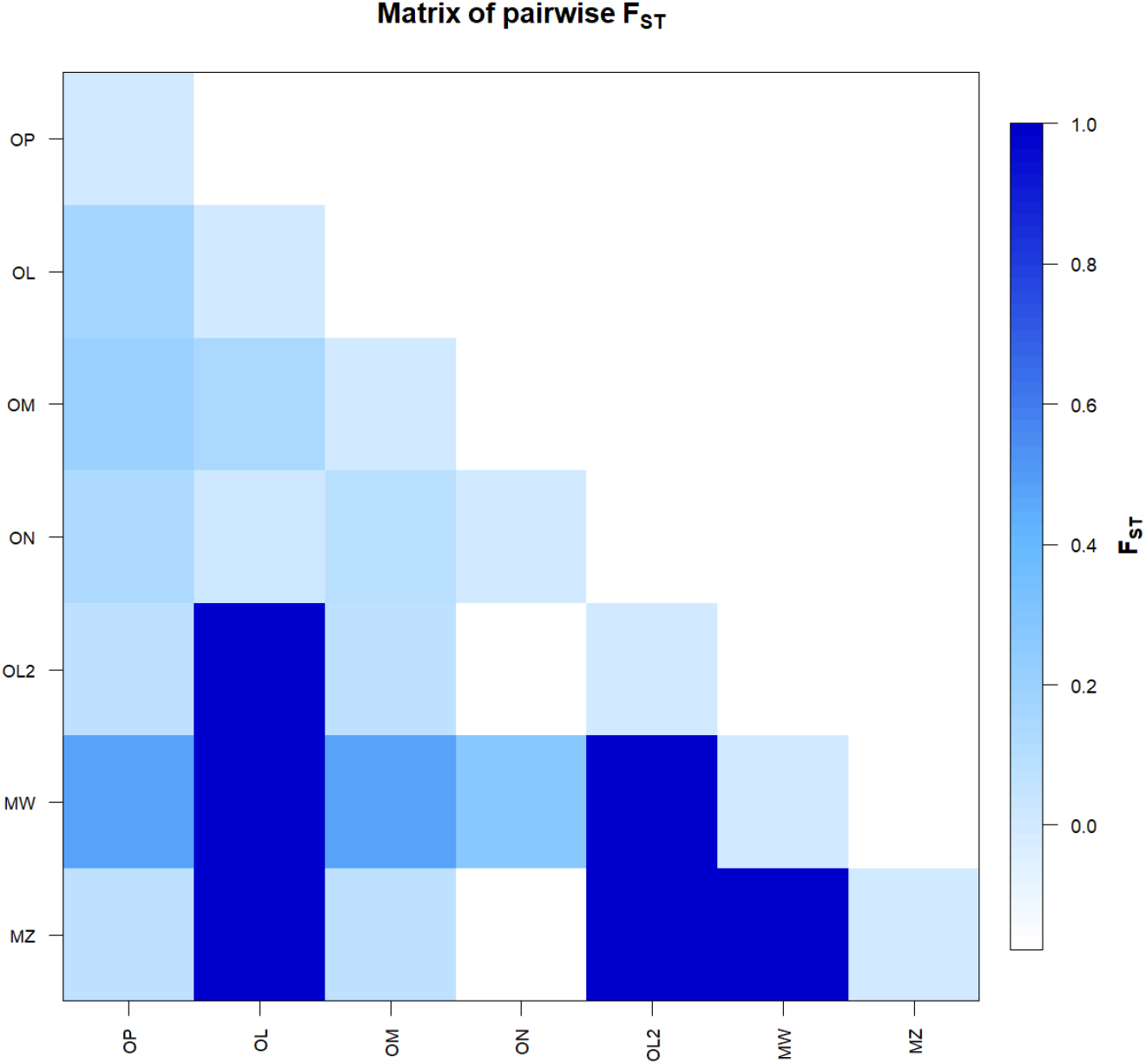
F_ST_-based genetic distance matrix between for the 28 sequences of the B.1.1 variant of SARS-CoV-2, from Brazilian states. * Generated by the statistical package in R language using the output data of the Software Arlequin version 3.5.1.2

**Figure 3.**
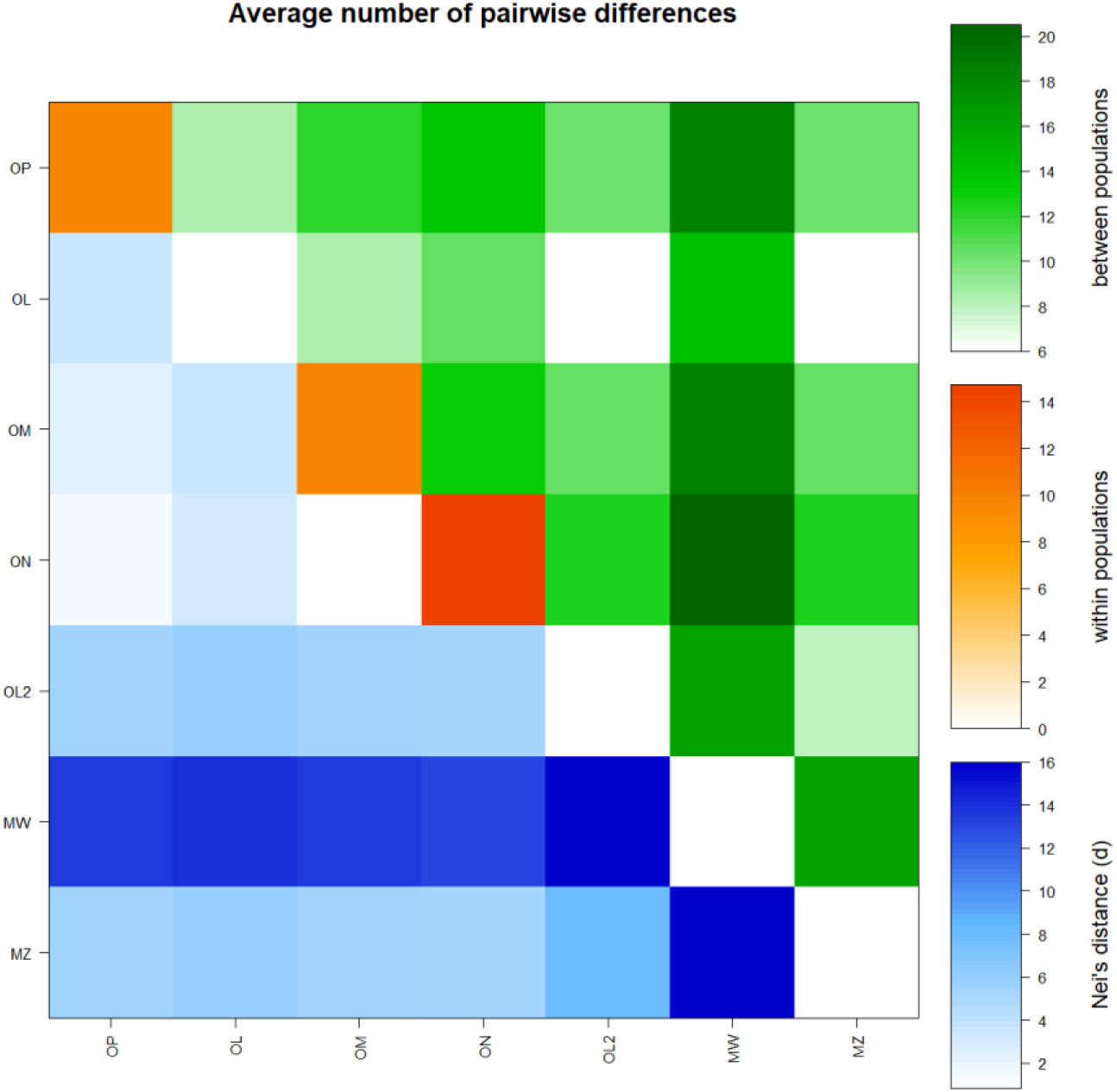
Matrix of paired differences between the populations studied: between the groups; within the groups; And Nei distance for the 28 sequences of the Variant B.1.1 of SARS-CoV-2, from Brazilian states.

**Table 2.** Demographic and spatial expansion simulations based on the τ, θ, and M indices of sequences for the 28 sequences of the variant B.1.1 of SARS-CoV-2, from Brazilian states.

| Statistics | OP | OL | OM | ON | OL2 | MW | MZ | Mean | s.d. |
| --- | --- | --- | --- | --- | --- | --- | --- | --- | --- |
| Demographic expansion |  |  |  |  |  |  |  |  |  |
| Tau | 12.65824 | 0.00000 | 6.99999 | 8.00000 | 0.00000 | 0.00000 | 0.00000 | 3.95118 | 5.22731 |
| Tau qt 2.5% | 5.18766 | 0.00000 | 5.19726 | 6.11328 | 0.00000 | 0.00000 | 0.00000 | 2.35689 | 2.95554 |
| Tau qt 5% | 5.51759 | 0.00000 | 6.02929 | 8.06445 | 0.00000 | 0.00000 | 0.00000 | 2.80162 | 3.57976 |
| Tau qt 95% | 18.33211 | 0.00000 | 12.63086 | 92.99963 | 0.00000 | 0.00000 | 0.00000 | 17.70894 | 34.03256 |
| Tau qt 97.5% | 19.30473 | 0.00000 | 13.87119 | 92.99963 | 0.00000 | 0.00000 | 0.00000 | 18.02508 | 34.00904 |
| Theta0 | 0.00352 | 0.00000 | 4.50000 | 19.89991 | 0.00000 | 0.00000 | 0.00000 | 3.48620 | 7.42946 |
| Theta0 qt 2.5% | 0.00000 | 0.00000 | 0.00000 | 0.00000 | 0.00000 | 0.00000 | 0.00000 | 0.00000 | 0.00000 |
| Theta0 qt 5% | 0.00000 | 0.00000 | 0.00000 | 0.00000 | 0.00000 | 0.00000 | 0.00000 | 0.00000 | 0.00000 |
| Theta0 qt 95% | 1.62599 | 0.00000 | 8.83124 | 96.39942 | 0.00000 | 0.00000 | 0.00000 | 15.26524 | 35.92188 |
| Theta0 qt 97.5% | 2.81959 | 0.00000 | 9.22851 | 96.39942 | 0.00000 | 0.00000 | 0.00000 | 15.49250 | 35.83722 |
| Theta1 | 23.02729 | 0.00000 | 3614.99145 | 3414.97837 | 0.00000 | 0.00000 | 0.00000 | 1007.57102 | 1713.88327 |
| Theta1 qt 2.5% | 6.28164 | 0.00000 | 38.32221 | 52.05584 | 0.00000 | 0.00000 | 0.00000 | 13.80853 | 21.92082 |
| Theta1 qt 5% | 8.38435 | 0.00000 | 50.52242 | 58.57455 | 0.00000 | 0.00000 | 0.00000 | 16.78305 | 26.08351 |
| Theta1 qt 95% | 186.77709 | 0.00000 | 727.50166 | 449.83169 | 0.00000 | 0.00000 | 0.00000 | 194.87292 | 288.86696 |
| Theta1 qt 97.5% | 213.65159 | 0.00000 | 864.99437 | 776.23733 | 0.00000 | 0.00000 | 0.00000 | 264.98333 | 388.34982 |
| SSD | 0.07292 | 0.00000 | 0.01398 | 0.06658 | 0.00000 | 0.00000 | 0.00000 | 0.02193 | 0.03312 |
| Model (SSD) p-value | 0.31000 | 0.00000 | 0.57000 | 0.02000 | 0.00000 | 0.00000 | 0.00000 | 0.12857 | 0.22572 |
| Raggedness index | 0.11338 | 0.00000 | 0.03160 | 0.08889 | 0.00000 | 0.00000 | 0.00000 | 0.03341 | 0.04820 |
| Raggedness p-value | 0.44000 | 0.00000 | 0.86000 | 0.96000 | 0.00000 | 0.00000 | 0.00000 | 0.32286 | 0.43304 |
| Spatial expansion |  |  |  |  |  |  |  |  |  |
| Tau | 9.19642 | 0.00000 | 7.64108 | 17.43719 | 0.00000 | 0.00000 | 0.00000 | 4.89639 | 6.82146 |
| Tau qt 2.5% | 4.34108 | 0.00000 | 3.52505 | 8.39432 | 0.00000 | 0.00000 | 0.00000 | 2.32292 | 3.26506 |
| Tau qt 5% | 4.86780 | 0.00000 | 4.76934 | 10.18609 | 0.00000 | 0.00000 | 0.00000 | 2.83189 | 3.95940 |
| Tau qt 95% | 15.21496 | 0.00000 | 11.39143 | 17.45551 | 0.00000 | 0.00000 | 0.00000 | 6.29456 | 8.04782 |
| Tau qt 97.5% | 17.27573 | 0.00000 | 11.69548 | 18.15338 | 0.00000 | 0.00000 | 0.00000 | 6.73208 | 8.63649 |
| Theta | 4.83152 | 0.00000 | 3.10712 | 0.01000 | 0.00000 | 0.00000 | 0.00000 | 1.13552 | 1.99883 |
| Theta qt 2.5% | 0.00072 | 0.00000 | 0.00072 | 0.00072 | 0.00000 | 0.00000 | 0.00000 | 0.00031 | 0.00039 |
| Theta qt 5% | 0.00072 | 0.00000 | 0.00629 | 0.00258 | 0.00000 | 0.00000 | 0.00000 | 0.00137 | 0.00237 |
| Theta qt 95% | 12.14673 | 0.00000 | 10.55732 | 7.95093 | 0.00000 | 0.00000 | 0.00000 | 4.37928 | 5.59718 |
| Theta qt 97.5% | 15.79439 | 0.00000 | 11.82491 | 9.28549 | 0.00000 | 0.00000 | 0.00000 | 5.27211 | 6.84282 |
| M | 138.78066 | 0.00000 | 9059.19716 | 4370.36352 | 0.00000 | 0.00000 | 0.00000 | 1938.33448 | 3532.90265 |
| M qt 2.5% | 8.83399 | 0.00000 | 39.24528 | 13.80161 | 0.00000 | 0.00000 | 0.00000 | 8.84013 | 14.50104 |
| M qt 5% | 12.94159 | 0.00000 | 44.80833 | 69.41346 | 0.00000 | 0.00000 | 0.00000 | 18.16620 | 27.93870 |
| M qt 95% | 1366.34285 | 0.00000 | 4336.44530 | 1808.30082 | 0.00000 | 0.00000 | 0.00000 | 1073.01271 | 1626.96592 |
| M qt 97.5% | 2104.32587 | 0.00000 | 5973.80135 | 4677.52366 | 0.00000 | 0.00000 | 0.00000 | 1822.23584 | 2541.32227 |
| SSD | 0.08379 | 0.00000 | 0.01393 | 0.03842 | 0.00000 | 0.00000 | 0.00000 | 0.01945 | 0.03174 |
| Model (SSD) p-value | 0.07000 | 0.00000 | 0.52000 | 0.41000 | 0.00000 | 0.00000 | 0.00000 | 0.14286 | 0.22381 |
| Raggedness index | 0.11338 | 0.00000 | 0.03160 | 0.08889 | 0.00000 | 0.00000 | 0.00000 | 0.03341 | 0.04820 |
| Raggedness p-value | 0.54000 | 0.00000 | 0.81000 | 0.74000 | 0.00000 | 0.00000 | 0.00000 | 0.29857 | 0.38107 |

### 4.3. Molecular diversity analyses

The molecular diversity analyses estimated by θ reflected a non-significant level of mutations among all haplotypes (transitions and transversions) and indel-type mutations (insertions or dissections) were not found in any of the studied groups (Table 3) (Figure 4). The D tests of Tajima and Fs de Fu did not show greater disagreements between the estimates of θ and general π, but with negative and highly significant values, indicating absence of population expansion (Table 4). The irregularity index (R= Raggedness) with parametric bootstrap simulated new θ values for before and after a supposed demographic expansion and in this case assumed a value equal to zero for the groups (Table 2).

**Figure 4.**
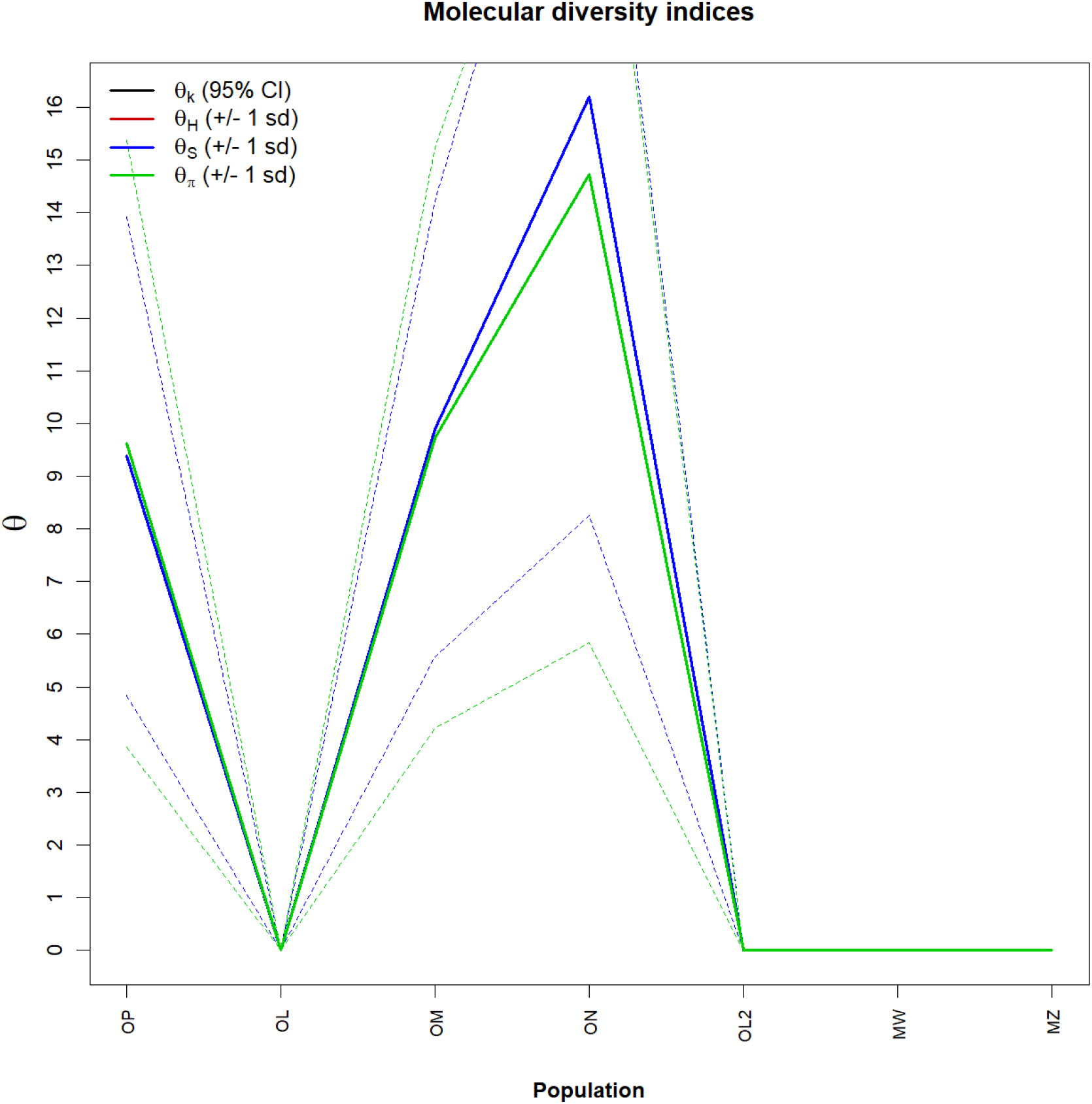
Graph of molecular diversity indices for the 28 sequences of the B.1.1 variant of SARS-CoV-2, from Brazilian states. In the graph the values of θ: (θk) Relationship between the expected number of alllos (k) and the sample size; (θH) Expected homozygosity in a balanced relationship between drift and mutation; (θS) Relationship between the number of segregating sites (S), sample size (n) and non-recombinant sites; (θπ) Relationship between the average number of paired differences (π) and θ. * Generated by the statistical package in R language using the output data of the Arlequin software version 3.5.1.2.

**Table 3.**
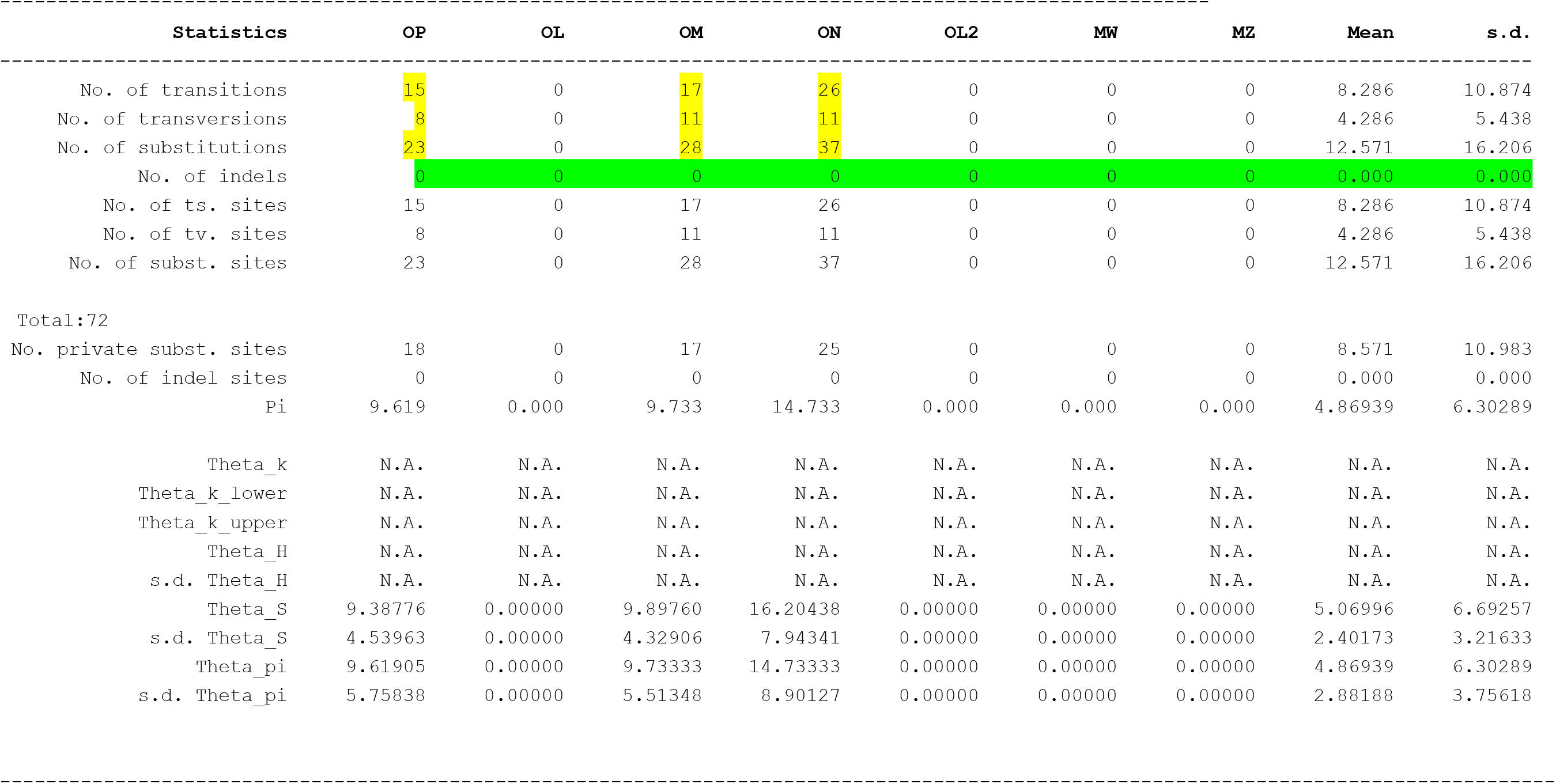
Molecular Diversity Indices for the 28 sequences of the Variant B.1.1 of SARS-CoV-2, from Brazilian states.

**Table 4.** Neutrality Tests for the 28 sequences of the Variant B.1.1 of SARS-CoV-2, from Brazilian states.

| Statistics | OP | OL | OM | ON | OL2 | MW | MZ | Mean | s.d. |
| --- | --- | --- | --- | --- | --- | --- | --- | --- | --- |
| **Ewens-Watterson test** |  |  |  |  |  |  |  |  |  |
| Sample size | 7 | 2 | 10 | 6 | 1 | 1 | 1 | 4.00000 | 3.65148 |
| No. of alleles(unchecked) | 7 | 2 | 10 | 6 | 1 | 1 | 1 | 4.00000 | 3.65148 |
| Observed F value | N.A. | N.A. | N.A. | N.A. | N.A. | N.A. | N.A. | N.A. | N.A. |
| Expected F value | N.A. | N.A. | N.A. | N.A. | N.A. | N.A. | N.A. | N.A. | N.A. |
| Watterson test: Pr(rand F <= obs F) | N.A. | N.A. | N.A. | N.A. | N.A. | N.A. | N.A. | N.A. | N.A. |
| Slatkin's exact test P-value | N.A. | N.A. | N.A. | N.A. | N.A. | N.A. | N.A. | N.A. | N.A. |
| **Chakraborty's test** |  |  |  |  |  |  |  |  |  |
| Sample size | 7 | 2 | 10 | 6 | 1 | 1 | 1 | 4.00000 | 3.65148 |
| No. of alleles(unchecked) | 7 | 2 | 10 | 6 | 1 | 1 | 1 | 4.00000 | 3.65148 |
| Obs. homozygosity | 0.00000 | 0.00000 | 0.00000 | 0.00000 | 0.00000 | 0.00000 | 0.00000 | 0.00000 | 0.00000 |
| Exp. no. of alleles | 5.47609 | 1.00000 | 7.13891 | 5.18084 | 0.00000 | 0.00000 | 0.00000 | 2.68512 | 3.11779 |
| P(k or more alleles) | N.A. | N.A. | N.A. | N.A. | N.A. | N.A. | N.A. | N.A. | N.A. |
| **Tajima's D test** |  |  |  |  |  |  |  |  |  |
| Sample size | 7 | 2 | 10 | 6 | 1 | 1 | 1 | 4.00000 | 3.65148 |
| S | 23 | 0 | 28 | 37 | 0 | 0 | 0 | 12.57143 | 16.20553 |
| Pi | 9.61905 | 0.00000 | 9.73333 | 14.73333 | 0.00000 | 0.00000 | 0.00000 | 4.86939 | 6.30289 |
| Tajima's D | 0.13919 | 0.00000 | -0.07945 | -0.57816 | 0.00000 | 0.00000 | 0.00000 | -0.07406 | 0.23150 |
| Tajima's D p-value | 0.57700 | 1.00000 | 0.50300 | 0.34300 | 1.00000 | 1.00000 | 1.00000 | 0.77471 | 0.28934 |
| **Fu's FS test** |  |  |  |  |  |  |  |  |  |
| No. of alleles(unchecked) | 7 | 2 | 10 | 6 | 1 | 1 | 1 | 4.00000 | 3.65148 |
| Theta_pi | 9.61905 | 0.00000 | 9.73333 | 14.73333 | 0.00000 | 0.00000 | 0.00000 | 4.86939 | 6.30289 |
| Exp. no. of alleles | 5.47609 | 1.00000 | 7.13891 | 5.18084 | 0.00000 | 0.00000 | 0.00000 | 2.68512 | 3.11779 |
| FS | -1.6316 | 0.00000 | -3.56086 | -0.39610 | 0.00000 | 0.00000 | 0.00000 | 0.000002 | 0.00000 |
| FS p-value | 0.10100 | 1.00000 | 0.02900 | 0.27500 | N.A. | N.A. | N.A. | N.A. | N.A. |

## 5. Discussion

With the use of phylogenetic analysis methodologies and population structure, it is not possible to detect the existence of similarity between haplotypes 28 sequences of variant B.1.1 of SARS-CoV-2, from the Brazilian states. Because significant levels of structuring were not found, we assumed that there are varying levels probably related to a gain of intermediate haplotypes over time, associated, perhaps, with a significant increase in gene flow. The non-occurrence of geographic isolations, because there are no past defragmentation events, may have generated this continuous pattern of genetic divergence between the groups, since the very low value found for genetic distance supports the absence of this pattern of divergence between haplotypes, as well as in the very low frequency of polymorphisms. This suggests that molecular diversity may be due to substitutions as the main components of variations. All analyses supportthe finding that there is a consensus in the conservation of the genome of the variant B.1.1 of SARS-CoV-2, from brazilian states, and therefore it is clearthat the genetic variability of mutants B.1.1. of the vírus of SARS-CoV-2 found in Brazil and used in the development of vaccines, reflects the efifid sciences of them parapleo minus this strain.

These considerations were also supported by simple phylogenetic pairing methodologies, such as UPGMA, which in this case, with a continuous pattern of genetic divergence between the groups, revealed a small number of branches with few mutational stages. These mutations were possibly not established by drift due to the founding effect, which accompanies the dispersive behavior and/or loss of intermediate haplotypes over the generations and the values found for the genetic distance considered the minimum differences between the groups, as well as the inference of values greater than or equal to those observed in the proportion of these permutations, including the *p-value of* the test.

The discrimination of the various genetic entities by distinct regions of the Brazilian territory was not perceived when the inter-haplotypic variations were hierarchised in all components of covariance: due to their intra- and inter-individual differences or their intra- and intergrupal differences, generating dendrograms that support the idea that there is no Significant differences should not be shared either in their form nor in their number, since the result of the estimates of the average evolutionary divergence for all groups was very low.

The φ estimators, although extremely sensitive to any form of molecular variation (Fu, 1997), supported the uniformity between the results found by all the methodologies employed, and can be interpreted as a phylogenetic confirmation that there is a consensus in the conservation of the sequences studied, and it is safe to affirm, iclusive, that the small number of found polymorphisms, reflit should godirectly into a low immune evasion. This is considered to ensure that the vaccines that used this strain are very efficient for the Brazilian population.

